# The sliding motility of the bacilliform virions of Influenza A Viruses

**DOI:** 10.1101/2023.03.21.533586

**Authors:** Laurie Stevens, Sophie de Buyl, Bortolo Matteo Mognetti

## Abstract

Influenza A virus (IAV) infection relies on the action of the hemagglutinin (HA) and neuraminidase (NA) membrane proteins. The HA ligands anchor the IAV virion to the cell’s surface by binding the sialic acid (SA) present on the host’s receptors while NA is an enzyme capable of cleaving the SA from the extracellular environment. It is believed that the activity of NA ligands increases the motility of the virions favoring the propagation of the infection. In this work, we develop a numerical framework to study the dynamics of a virion moving across the cell surface for timescales much bigger than the typical ligand-receptor reaction times. We find that the rates controlling the ligand-receptor reactions and the maximal distance at which a pair of ligand-receptor molecules can interact greatly affect the motility of the virions. We also report on how different ways of organizing the two types of ligands on the virions’ surface result in different types of motion that we rationalize using general principles. In particular, we show how the emerging motility of the virion is less sensitive to the rate controlling the enzymatic activity when NA ligands are clustered. These results help to assess how variations in the biochemical properties of the ligand–receptor interactions (as observed across different IAV subtypes) affect the dynamics of the virions at the cell surface.

## I. INTRODUCTION

Influenza A Viruses (IAVs) are pathogens that infect cells in the respiratory tract and cause seasonal epidemics and flu pandemics. Its virions resemble spherical or filamentous particles,^1^ the latter with a diameter between 70 nm and 90 nm and an axial length between 100 nm and 50 *μ*m.^2,3^ Bacilliform virions are filamentous particles with an axial length smaller than 250 nm are believed to be the vectors spreading the infection from cell to cell.^3–5^ IAVs are classified based on the possible subtypes of hemagglutinin (HA) and neuraminidase (NA) membrane proteins (in the following ligands) found at the surface of the virus particle (Fig. 1*a*).^2,6,7^ HA ligands binds the sialic acid (SA) present as the end-group of both the glycoproteins at the cell surface (in the following receptors) and the sugar chains found in the extracellular medium (constituted by mucin strands)^8–12^. Instead, NA molecules are enzymes that cleave the SA molecules,^13^ therefore limiting the number of HA–SA contacts (in the following bridges) between the host and the pathogen.^2,14^

**FIG. 1.**
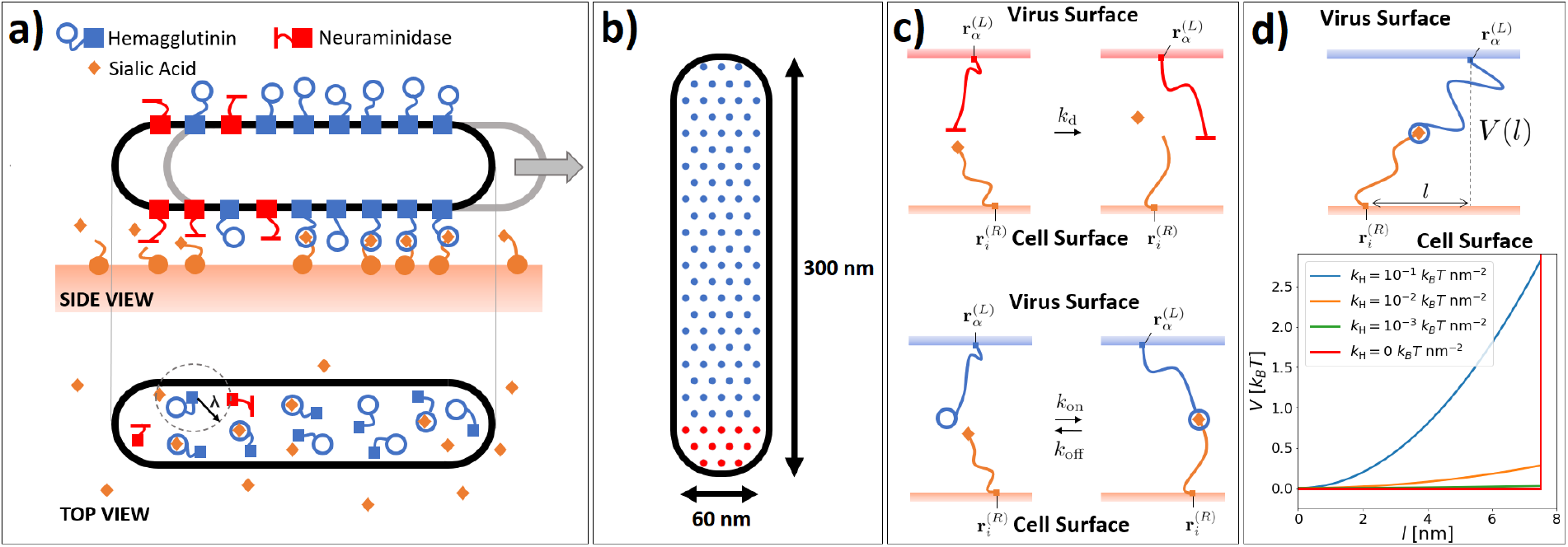
(*a*) Bacilliform IAV particles carry Hemagglutinin (HA) and Neuraminidase (NA) ligands, respectively, binding and cleaving the sialic acid (SA) presented by receptor molecules. (*b*) 2D representation of a bacilliform virion.^16^ The IAV is mapped into a 2D spherocylinder (corresponding to the top view of Fig. 1*a* bottom) with ligands distributed as in a triangular lattice. (*c*) Possible ligand-receptor reactions and corresponding rates: (*top*) NA ligands passivate receptors with a rate *k*_d_; (*bottom*) HA ligands bind/unbind receptors carrying SA with a rate 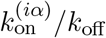. Ligand-receptor pairs react only if the distance between their tethering points 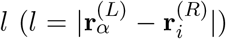 is smaller than *λ (d) (top)* A bridge exerts a force on the virion due to a harmonic potential *V* (*l*), *V* (*l*) = *k*_*H*_*l*^2^*/*2; (bottom) Harmonic potentials considered in this work for *λ* = 7.5 nm.

It is now accepted that successful infection of an IAV strand requires a synergic activity between NA and HA ligands.^15–17^ For instance, the process of exposing IAV to NA inhibitors generates IAV mutants with reduced HA-SA affinity.^18^ The implications of the synergy between HA and NA ligands on the host-pathogen dynamics is a matter of debate. It is believed that NA ligands help the virus to navigate the extracellular medium by reducing the number of HA-SA bridges and generating local gradients in SA concentration.^13,19,20^ The number of bridges and SA gradients are expected to affect the motility of the pathogen as characterized in theoretical studies of synthetic walkers.^21–24^ Mobile viruses are likely more infectious, given that they do not remain trapped in the extracellular matrix and can navigate the cell surface to find a spot favorable to endocytosis.^13,25^

This work aims at predicting the emerging motility of IAV particles using a coarse-grained model in which a single bacilliform virion moves along a surface carrying receptors tipped by SA. We validate an efficient and quantitative reaction–diffusion simulation protocol allowing us to generate thousands of 100 s–long trajectories. We then characterize the virion motility as a function of the model’s parameters, namely, the rates controlling the reactions between ligand-receptor (*k*_on_, *k*_off_, and *k*_d_), the maximal distance at which a pair of ligand–receptor molecules can interact (in the following defined as the ‘interaction range’, *λ*), the receptors’ diffusion constant (*D*_R_), and the ligands’ organization at the virus surface. We mainly consider viruses in which NA and HA ligands are segregated. At the simulation timescales (∼ 100 s), the pathogen then follows a diffusive motion with a drifting component, the latter due to the dynamic gradient in SA concentration. We extensively use the drifting velocity (*v*_*y*_) to characterize the motility of the virus particle as a function of the system parameters. We find that *v*_*y*_ is mainly affected by the number of bridges (controlled by the rates at which bridges are formed/destroyed) and the bridges’ maximal extensibility (*λ*). The rate controlling the catalytic activity of the NA (*k*_d_) can affect the drifting of the virions at low numbers of bridges. As shown in Ref.^16^, we find that *v*_*y*_ is maximized at finite values of the receptor diffusion constant (*D*_R_). Finally, we consider systems in which NA and HA ligands are uniformly distributed over the virus’ surface. Similarly to what was reported in models of Influenza C viruses^26^ or in similar synthetic walkers,^27^ we show how in this case the direction of motion is rotated by 90°, with the virus particles moving preferentially orthogonally to its axis. Moreover, for uniform NA distributions, the virion motility is more sensitive to *k*_d_.

This paper contributes to establishing a quantitative link between molecular parameters and the emerging motility of the pathogen. Such a link allows correlating the motility with the infectivity of different IAV subtypes. Our results also extend the rich literature on synthetic molecular walkers (e.g. using burnt-bridge schemes^27–33^) and suggest new designs featuring tailored types of motion.

## II. THE MODEL

### A. Model description

Being interested in the sliding motion of the pathogen, we build upon the 2D model recently introduced by Vahey and Fletcher^16^ in which the distance between the virus and the surface is constant. The use of a 2D model is justified by the fact that the particles never detach from the surface as we never recorded configurations in which *n*_*b*_ = 0. We map the virion into a 2D spherocylinder (corresponding to the projection of the 3D particle onto the surface, Fig. 1*a*), with a length (*L*) and width (*d*) equal to *L* = 300 nm and *d* = 60 nm (Fig. 1*b*). Ligands are distributed over the 2D particle and are taken at a fixed distance from the surface plane (Fig. 1*b, c, d*). Without ligand-receptor bridges, the particle diffuses with translational and rotational diffusion constants equal to (*D*_‖_, *D*_⊥;_) and *D*_*θ*_, respectively. *D*_‖_ and *D*_⊥;_ are, respectively, the diffusion constant in the parallel and perpendicular direction with respect to the particle’s axis. On the surface of the particle, the ligands are distributed like in a triangular lattice with a unit length equal to 15 nm (see Fig.1*b*).^16^ Given that the stoichiometric ratio between HA and NA ligands is equal to [HA]/[NA]= 5*/*1, the virus carries 97 (12) HA (NA) ligands. With the exceptions of the systems studied in Sec. V, we consider ‘polarized’ particles in which the two types of ligands are segregated and occupy the tail (NA) and head (NA) of the virion (Fig. 1*b*). The surface is decorated by uniformly distributed receptors with density *ρ*_R_. Receptors are fixed except for the systems considered in Sec. IV C where they follow a Brownian motion with a diffusion constant equal to *D*_R_.

Each receptor can be found in three different states: bound to a ligand, unbound and active (i.e., carrying a SA molecule), and inactive (unbound without SA). A ligand–receptor pair can form a bridge only if the lateral distance between the tethering points (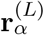 and 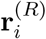) is smaller than the interaction range, *λ*, 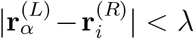 (Fig. 1*d*). Accordingly, bridges limit the configuration space available to the virus. The force acting on the virus because of a bridge is given by 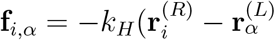 where *k*_*H*_ is a harmonic constant (Fig. 1*d*).

We consider three different types of reaction, and therefore three associated reaction rates (Fig. 1*c*): 1) An active, unbound receptor (*i*) in the reach of an unbound HA ligand (*α*) can form a bridge with a rate equal to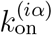; 2) A bridge can be opened with a rate *k*_off_, leading to the formation of an unbound HA ligand and an unbound, active receptor; 3) An NA ligand (*α*) can inactivate a receptor (*i*) with a rate *k*_d_ (if 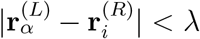). Reaction 3) prevents the desialylated receptor from taking part in reaction 1).

The formation of a bridge alters the configuration energy of the system as it adds a harmonic spring tethering the virus particle to the surface (see Fig.1*d*). Detailed balance conditions then require that the *on* and/or *off* rates be configuration dependent as follows

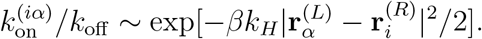

In the following, we consider configuration–dependent *on* rates and constant *off* rates. Note that the typical values of the forces *f* considered in this work (*f* ≤ 2.7 pN, see Fig. 1*d*) are 5 times smaller than the equilibrium force *f*_eq_ of the NA–HA system (*f*_eq_ = 15.1 ± 0.8 pN^34^), where *f*_eq_ is the transition point from the equilibrium to the kinetic regime in pulling experiments.^35,36^ We then write the *on* rate 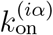 as follows

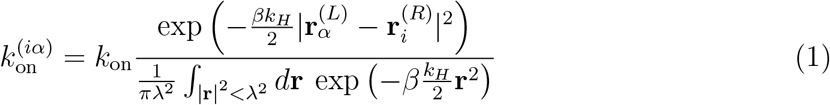

where the denominator of the r.h.s. term is equal to

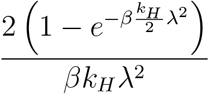

The use of such a normalization in the r.h.s. of Eq. 1 may look awkward as compared to standard expressions.^37^ However, the use of Eq. 1 guarantees that the average number of HA-SA bridges (⟨*n*_*b*_⟩) is not affected by *k*_*H*_. Given that, as discussed in Sec. IV A, changes in ⟨*n*_*b*_⟩ have a major impact on the motility of the virion, Eq. 1 allows assessing the effect of different *k*_*H*_ on the motility of the particle by using a single value of *k*_on_ and *k*_off_. *k*_on_ is the *on* rate for *k*_*H*_ = 0 and is directly linked to experimental values in the next section.

### B. The simulation parameters

We first discuss the reaction rates. These are measured in experiments tracking the reaction kinetics between SA molecules and free (i.e., not supported by a virus particle) HA/NA molecules.^17,34,38^ These studies set the *on* rate (*k*_on,0_) leading to the formation of an HA-SA complex to *k*_on,0_ ≈ 150 − 1000 M^−1^s^−1^. From *k*_on,0_, we derive an estimation of *k*_on_ (Eq. 1) by modeling grafted HA and SA molecules as free complexes confined in a sphere with a volume equal to 4*πλ*^3^*/*3.^16^ In particular

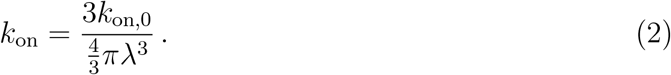

The previous expression includes an avidity term arising from the trimeric nature of the hemagglutinin (featuring three different sites capable of docking SA molecules). Note that, in our current model, two or more SA receptors cannot bind to the same HA ligand. Using *k*_on,0_ = 200 M^−1^s^−1^ and *λ* = 7.5 nm in Eq. 2 gives *k*_on_ = 0.56 s^−1^. This is the value that has been used to obtain most of the results presented in this paper (for *λ* = 7.5 nm). The *off* rate (*k*_off_, Fig. 1*c*) has been chosen as a tuning parameter resulting in a sought number of bridges. The range of values considered (see Tab. I) are consistent with experimental results (e.g., Ref.^34^ reports *k*_off_ = 66 ± 13 s^−1^).

**TABLE I.** Simulation parameters employed by this study as justified in Sec. II

| Parameters | Value Range |
| --- | --- |
| **Virus and receptors** |  |
| $L$ | 300 nm |
| $d$ | 60 nm |
| $\rho_R$ | $0.02 - 0.2 \text{ nm}^{-2}$ |
| $k_H$ | $10^{-3} - 10^{-1} k_B T \text{ nm}^{-2}$ |
| $T$ | 308 K |
| **Reactions** |  |
| $k_{\text{on}}$ | $0.12 - 2.0 \text{ s}^{-1}$ |
| $k_{\text{off}}$ | $4.0 - 75.0 \text{ s}^{-1}$ |
| $k_d$ | $0.09 - 30000 \text{ s}^{-1}$ |
| $\lambda$ | $3.75 - 15.0 \text{ nm}$ |
| **Diffusion** |  |
| $D_{\parallel}$ | $3.38 \cdot 10^6 \text{ nm}^2/\text{s}$ |
| $D_{\perp}$ | $1.75 \cdot 10^6 \text{ nm}^2/\text{s}$ |
| $D_{\theta}$ | $2.75 \cdot 10^2 \text{ rad/s}$ |
| $D_R$ | $10^{-5} - 10^5 \text{ nm}^2/\text{s}$ |

The desialylation of a receptor carrying a SA group by an NA ligand is described by the following chemical equation

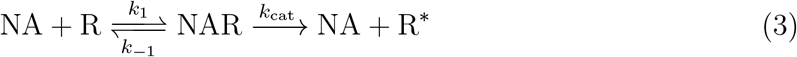

where NAR is a transitory bridge formed by a receptor (R) and the NA ligand. R and R^*^ denote, respectively, a receptor with and without SA. Experiments similar to the ones discussed above set *k*_1_ to a value comparable with *k*_on_.^34^ On the other hand, the measured values of *k*_cat_ are usually between 10 s^−1^ and 100 s^−1^, much bigger than *k*_1_ and *k*_−1_.^17,34,38^ As done in Ref.^16^, we then neglect the intermediate state and approximate the desialylation process as a bimolecular reaction with a rate (*k*_d_) equal to *k*_d_ = *k*_1_. We note that existing literature often employed much bigger values of *k*_d_ neglecting that, if *k*_cat_ → ∞, *k*_d_ is diffusion limited. To facilitate the comparison with other models, in this work we also consider *k*_d_ as a tuning parameter scanning multiple orders of magnitude. Variations in the values of the reaction rates are also justified by the existence of multiple families of HA/NA molecules and the fact that ligands can recognize different types of glycans with different affinities.^39–42^

The motion of a 2D spherocylinder is described by the diffusion constant of the particle moving along and perpendicularly to its axis (respectively, *D*_‖_ and *D*_⊥;_), and the rotation diffusion constant (*D*_*θ*_). We employ the values used in Ref.^16^ as obtained using Brenner’s theory.^43^ In the next section, we show how the emergent motion of the particle is solely determined by the reaction dynamics (with *D*_‖_, *D*_⊥;_, and *D*_*θ*_ playing no role). The timescale of the dynamics is then fully determined by the reaction rates (*k*_on_, *k*_off_, and *k*_d_).

Tab. I summarises all the parameters used in the present paper. The simulation programs employed to generate the results of this work are available at the following link.^44^

## III. SIMULATING THE REACTION-DIFFUSION DYNAMICS

### A. Simulation chart flow

The model presented in Sec. II is simulated using a reaction-diffusion algorithm^24,45–48^ whose chart flow reads as follows

1. *t* ← 0
2. **while** *t < t*_*E*_ **do**
3. *Reaction*(Δ*t*)
4. *Diffusion*(Δ*t*)
5. *t* ← *t* + Δ*t*
6. **end while**

where *t* is the time variable, Δ*t* the simulation timestep, and *t*_*E*_ the time at the end of the simulation. *Reaction*(Δ*t*) implements ligands/receptors reactions, while *Diffusion*(Δ*t*) updates the particle’s and the receptors’ position (when mobile) for a given set of HA–SA linkages. For the reaction part, we use the Gillespie algorithm^49^ adapted to the case in which each ligand/receptor is treated as a different chemical species. This complication arises from the fact that the *on* rates are configuration dependent (see Eq.1) and from the fact that different reactions of the same type constrain the particle’s motion differently. In particular, our algorithm not only evolves the number of bridges but also tracks the list of ligand/receptor molecules that form a bridge.

Reaction(Δ*t*) samples the list of reactions and the time for them to happen (*τ, J* =1, 2,*·*··). The list of reactions is terminated just before the cumulative reaction time exceeds Δ*t*, 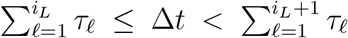, where *i*_*L*_ labels the last reaction implemented by *Reaction*. The choice of the reaction is made by sampling time–dependent affinity matrices *a*^X^[*α*][*i*] defined as the rate at which ligand *α* and receptor *i* undergo a reaction of type *X* (*X* = *on/off* for the formation/breakage of a bridge, *X* = *d* for a desialylation event). In particular, 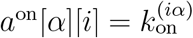 if αand *i* can form a bridge (*a*^on^[*α*][*i*] = 0 otherwise), *a*^off^ [α][*i*] = *k*_off_ if *α* and *i* form a bridge, and *a*^d^[α][*i*] = *k*_d_ if αcan inactivate *i*. Given *a*^X^[α][*i*], we define *a*^X^[α] (*a*^X^[α] = *a*^X^[*α*][*i*]), 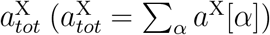, and 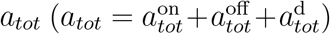. A reaction is then selected by following the decision tree of Fig. 2 in which we first choose the type of reaction *X* (with probability 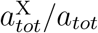), then the ligand αinvolved in the reaction (with probability 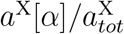), and finally the receptor *i* (with probability *a*^X^[*α*][*i*]*/a*^X^[*α*]). Given *a*_*tot*_, we sample *τ ℓ* from a Poisson distribution with average ⟨*τ*_*ℓ*_⟩= (*a*_*tot*_)^−1^. Before sampling the next reaction, we update the affinity matrices (along with *a*_*tot*_, 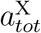, and *a*^X^[α]) using Verlet lists.

**FIG. 2.**
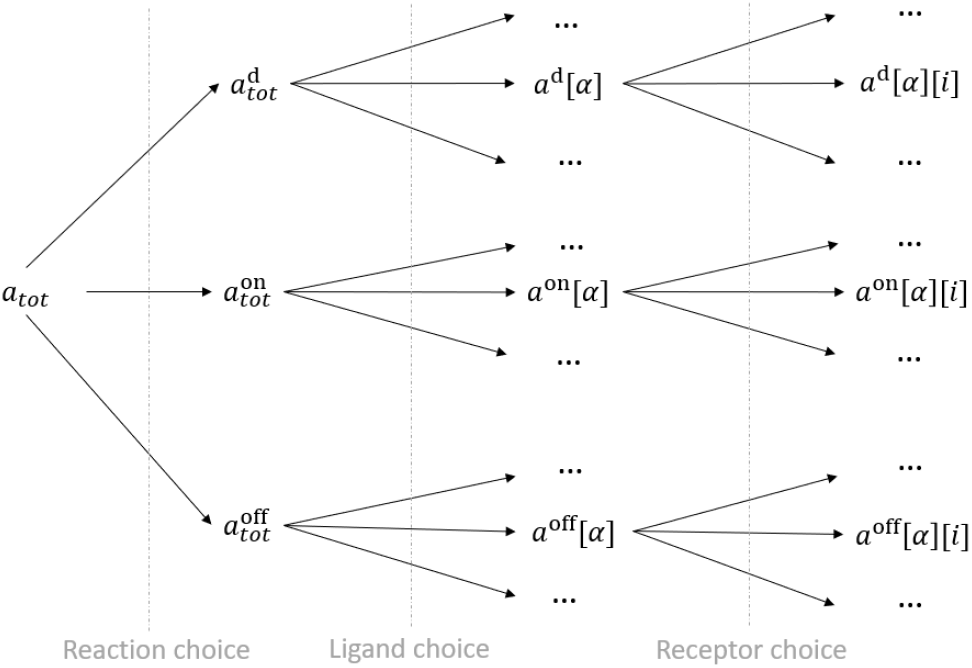
Decision tree employed in the selection of the reacting molecules (*α* and *i*) and the type of reaction (*d, on*, or *off*).

For the diffusion update, *Diffusion*(Δ*t*), either use Brownian dynamics updates (below) or an equilibrium sampling of the configuration space available to the particle for a given set of bridges (Sec. III B). When using Brownian dynamics simulations, the center of mass (*x, y*) and the orientation *θ* of the particle are updated as follows:

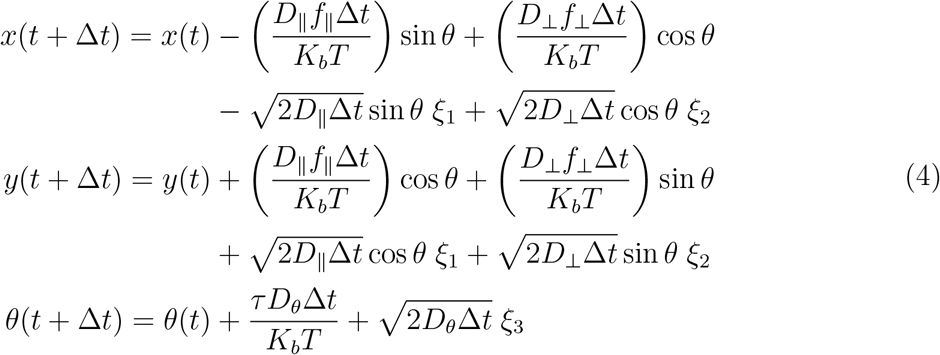

where *f*_‖_ and *f*_⊥;_ are the components of the total force (obtained by summing over all bridge contributions, **f**_*i, α*_, see Sec. II) in the virion reference frame

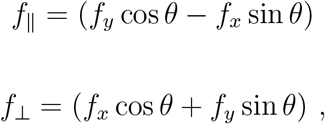

*τ* is the torque, and *ξ*_*i*_ are random numbers following the standard normal distribution. The Brownian update(Eq. 4) is rejected if, for the proposed new configuration, an existing bridge becomes overstretched. A high number of bridges sensibly reduces the particle configuration volume (see Fig. 3)^24^ leading to a high rejection rate. High rejection rates can slow down the dynamics in a way that depends on the simulation parameters (in particular Δ*t*). To reduce the number of rejections and avoid biases, we divide the Brownian step update into *N*_*B*_ substeps evolving the particle’s coordinates by a time step equal to Δ*t/N*_*B*_. *N*_*B*_ is calculated by constraining the typical Brownian update to remain smaller than the expected size of the configuration area available to the particle at a given orientation (Ω in Fig. 3c). However, in typical simulations, this choice may result in a prohibitively large number of sub-steps (up to *N*_*B*_ = 5· 10^5^). Therefore, in the next section, we elaborate on a more efficient method in which the configuration of the particle is taken as uniformly distributed in Ω.

**FIG. 3.**
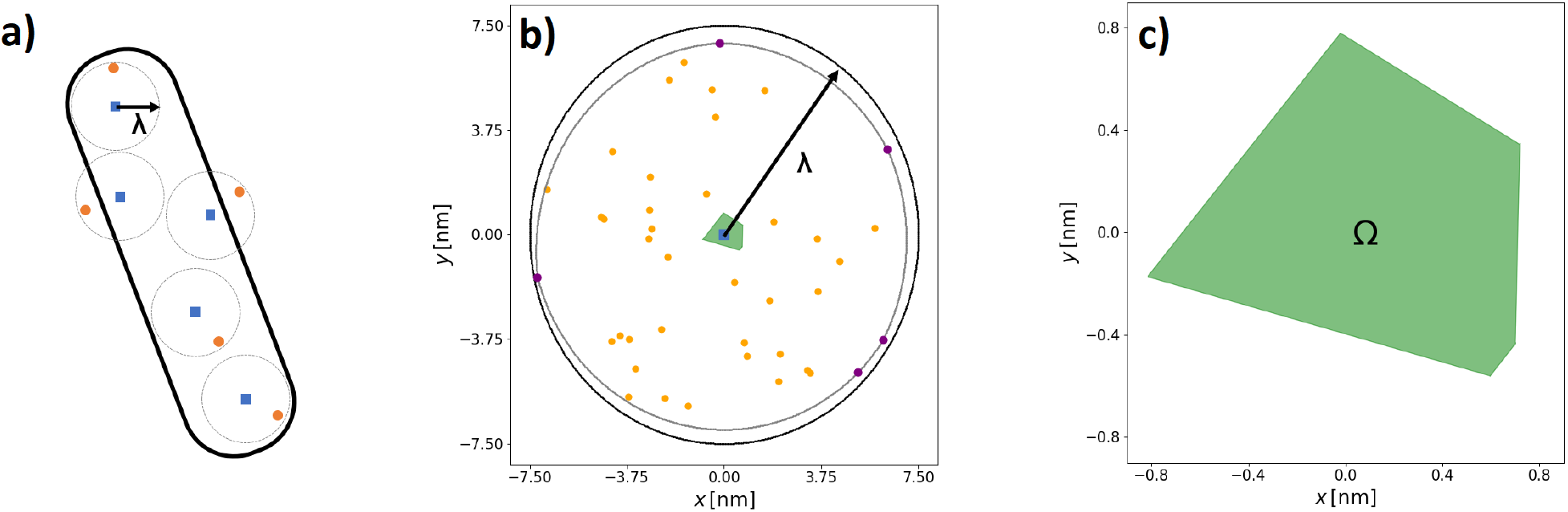
(*a*) The possible positions and orientations of the virion are determined by the bridges as the ligand-receptor distance is constrained to remain smaller than *λ*. (*b*) Geometrical construction employed in Ref.^24^ in which each pair of bound molecules (represented by squares and circles in *a*) is rigidly translated to overlap all ligands’ position (squares) with the center of mass of the particle. The dynamics of the system (at a given *θ*) are then described by a circle of radius *λ* rattling around the receptors (circles). The *n*_*cb*_ constraining bridges correspond to the receptors that can enter into direct contact with the border of the rattling circle (purple points). For clearness, in *a* we use fewer bridges than in *b* (in *b* we have *n*_*b*_ = 40 as in most of the systems studied in this work). (*c*) Ω is the configuration space scanned by the center of mass in the rattling dynamics. Ω is constituted by a number of vertices equal to *n*_*cb*_ jointed by arcs of radius *λ*.^24^ For 40 bridges and *λ* = 7.5 nm, the size of Ω is ≈ 1 nm.

### B. Equilibrium update of the virion’s coordinates

As investigated in our previous study,^24^ at short time scales, the dynamics of the system can be described in terms of a diffusive motion in which the system explores the finite configuration space available to the center of mass and orientation of the particle at a given set of bridges (Ω in Fig. 3*c*). At larger time scales, the dynamics are determined by the evolution of the particle’s configuration space due to the reconfiguration of ligand–receptor bridges. Importantly, only a subset of bridges (in the following labeled by constraining bridges, Fig. 3*b*) determines the particle’s configuration space. In particular, at a given particle orientation (*θ*), the center of mass of the particle can explore a 2D polygon with curved edges (Ω in Fig. 3*c*).^24^ The *n*_*cb*_ edges of Ω(*θ*) correspond to the particle’s configurations in which a constraining bridge is maximally stretched (Fig. 3*b*). In Ref.^24^, we showed that *n*_*cb*_ remains finite (even when *n*_b_ → ∞) and is well approximated by *n*_*cb*_ ≈ 5. It follows that the average time (*τ*_*r*_) taken by a reaction (e.g., the breakage of a constraining bridge) to update Ω reads as *τ*_*r*_ = (2*n*_*cb*_*k*_off_)^−1^.^24^ On the other hand, the time (*τ*_*d*_) taken for the particle to diffuse over a distance comparable with the size of Ω (≈ 1 nm when *n*_*b*_ = 40, Fig. 3c) is 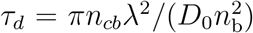, ^24^ where *D*_0_ is the typical diffusion constant of the particle when free from bridges (*D*_0_ ∼ *D*_‖_, *D*_⊥_ see Tab. I). The fact that *τ*_*r*_ *≫τ*_*d*_ (*τ*_*r*_*/τ*_*d*_ *>* 10^4^, see Tab. I) justifies decomposing the dynamics into a short-time subdiffusive motion (in which the particle rattles inside Ω) and a slow drifting motion due to the evolution of Ω.

Based on the previous considerations, in this work, we tested a coarse-grained simulation algorithm that simplifies the rattling dynamics by substituting the Brownian updates (which are computationally demanding given the prohibitively large numbers of sub-steps *N*_*B*_, Sec. III) with an equilibrium sampling of the configuration space for a given set of bridges. The new position of the center of mass of the particle, (*x*(*t* + Δ*t*), *y*(*t* + Δ*t*)), is chosen with uniform probability inside Ω. After updating the center of mass, we determine the range of possible orientations compatible with the existing bridges, Δ*θ*, and sample the new orientation inside Δ*θ*. Importantly, as compared to Brownian dynamics updates (Sec. III), this method allows us to speed up the simulations by a factor of 10.

As we sample the center of mass position uniformly in Ω (similarly for the orientation), we have that our approach is strictly consistent only with a microscopic model in which the bridges do not exert any force on the particle (*k*_*H*_ = 0 in Fig. 1*d*). To assess the impact of the bridging force on the motility of the particle, in Fig. 4 we report the sampled drifting velocity (*v*_*y*_), calculated as detailed in the next section, for different harmonic constants *k*_*H*_ by employing the Brownian dynamics updates (Eqs. 4). Fig. 4 shows that the bridging force has no statistical impact on *v*_*y*_. This is expected given that bridging forces affect the rattling dynamics which is not relevant to the dynamics at long time scales. Moreover, the predictions of *v*_*y*_ at different *k*_*H*_ are consistent with the value obtained using the coarse-grained algorithm described above (dashed line) validating the proposed method. The stability of the simulation method when changing the integration time step (Δ*t*) is proven in ESI Tab. 1.

**FIG. 4.**
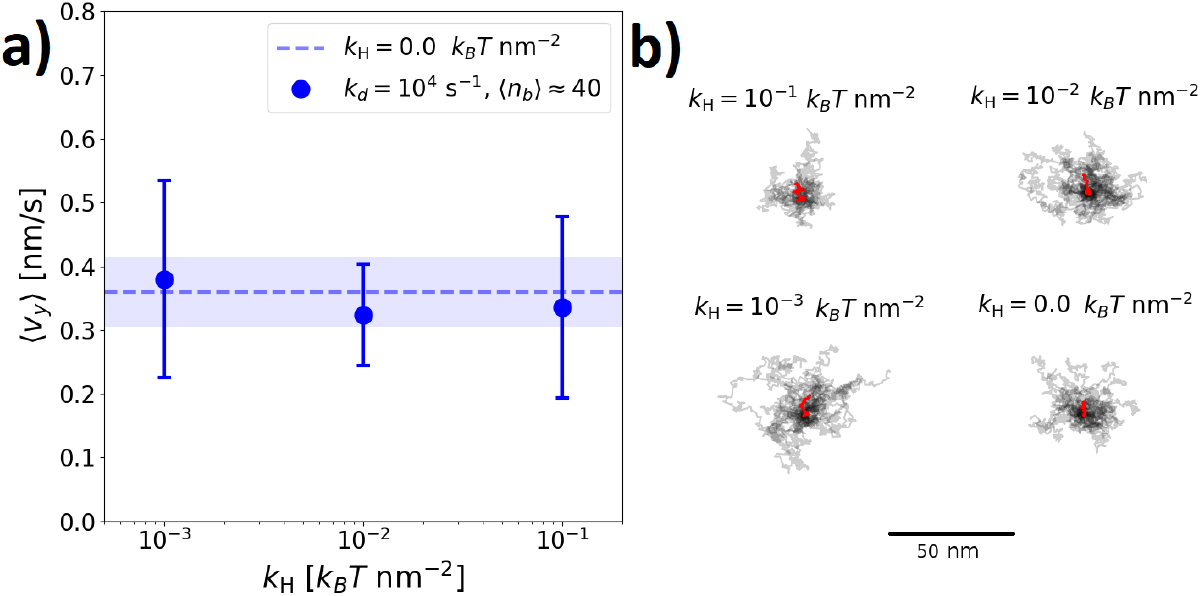
(*a*) Drifting velocities (Sec. IV) and (*b*) center of mass trajectories of systems employing different bridging potentials (Fig. 1*d*). In (*b*) we report 20 trajectories (highlighted in red is the averaged one). The bridging potential has no impact on the long time dynamics of the system. We use *k*_off_ = 25.0 s^−1^, *k*_on_ = 0.56 s^−1^, *k*_d_ = 10^4^s^−1^ (resulting in an average number of bridges equal to ⟨*n*_b_ ⟩ = 40), *ρ*_R_ = 0.2 nm^−2^, Δ*t* = 10^−3^ s, and *t*_*E*_ = 100 s.

## IV. DRIFTING VELOCITY *v*_*y*_

Fig. 5a reports the simulation setting employed in this work. We prepare a surface functionalized by receptors carrying sialic acid (in orange) and place the particle in the middle of the surface pointing towards the positive direction of the *y* axis. We then evolve the system for *t*_*E*_ = 100 s using the coarse-grained dynamics described in Sec. III. Fig. 5a shows how, for *t >* 0, patches depleted from sialic acid readily form. The motility of the particle is sufficiently slow to allow the NA ligands to remove all the sialic acid found in the region of the surface scanned by the tail of the particle. The region depleted from sialic acid acts as a damp pushing the particle towards the positive direction of the *y* axis (see Sec. IV B and Sec. V for further explanations on the driving force pushing the particle away from the SA depleted region). This is shown in Fig. 5 where we report the *y* coordinate of the center of mass as a function of time for 30 trajectories (gray lines). The averaged trajectory (red line) features a linear trend from which we extract the drifting velocity *v*_*y*_ (blue line). For very long trajectories, we do not exclude that the particle may lose persistency resulting in smaller values of *v*_*y*_. To have affordable simulations and significant statistics, in this work we assess the degree of motility of polarized virions using the drifting velocity calculated as in Fig. 5.

**FIG. 5.**
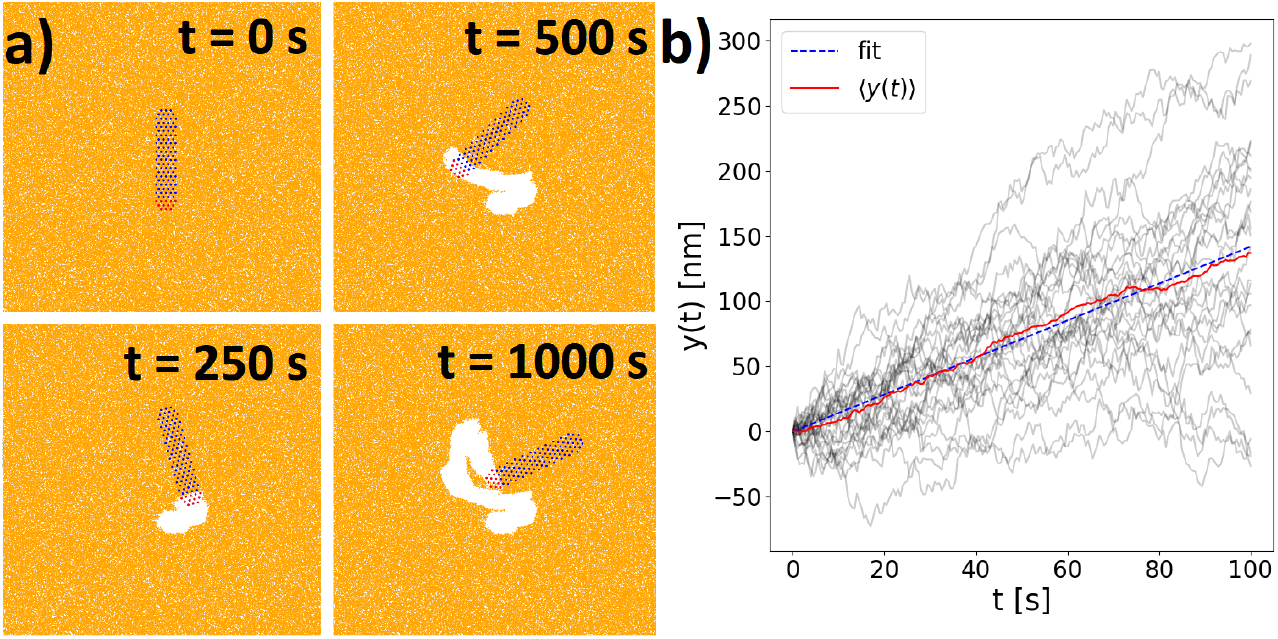
(*a*) Simulation snapshots at different times *t*. The region depleted from SA (in white) pushes the particles toward the positive direction of the *y* axis. A movie showing the evolution of the full trajectory is available in the ESI. (*b*) Position of the *y* coordinate of the particle’s center of mass as a function of *t*. We report 30 trajectories (gray lines), the average trajectory (red line), and the fitted trajectory (dashed, blue line) from which we extract the drifting velocity *v*_*y*_. We use *k*_off_ = 25.0 s^−1^, *k*_on_ = 0.56 s^−1^, *k*_d_ = 10^4^ s^−1^, ⟨*n*_b_ ⟩ = 20, *ρ*_R_ = 0.2 nm^−2^, and Δ*t* = 10^−3^ s.

### A. Number of bridges *n*_b_ and desialysation rate *k*_d_

In this section, we study the impact of *k*_d_ and *n*_b_ on the drifting velocity *v*_*y*_, the latter calculated as described in Fig. 5. As discussed in Sec. III B, the emerging dynamics is controlled by the rate at which the configuration space available to the particle’s center of mass (Ω) is updated by a reaction, (*τ*_*r*_)^−1^ ∼ *k*_off_.^24^ On the other hand, the results of Ref.^24^ show how, following an updated of Ω, the center of mass of the particle moves, on the average, of a quantity *δ* with *δ* ∼ ⟨*n*_b_ ⟩^−1^. These considerations allowed us to derive the following scaling law for the diffusion constant *D* of passive particles (i.e., particles not carrying NA ligands)^24^

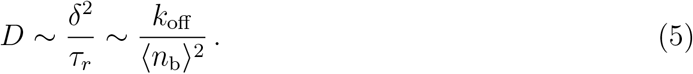

In the presence of NA ligands, we expect the region depleted from sialic acid to bias the motion of the particle towards positive values of the *y* axis (see Fig. 5a). We then expect the drifting velocity to scale like ⟨*v*_*y*_ ⟩ ∼ *δ/τ*_*r*_ ∼ *k*_off_ */* ⟨*n*_b_ ⟩.

We test and complement this scaling law using the simulation results of Fig. 6*a*. In these simulations, we keep *k*_on_ constant while changing *k*_off_ and *k*_d_ (with *k*_on_*/k*_d_ constant). Fig. 6a confirms the trend of the scaling law for ⟨*v*_*y*_ ⟩ proposed above. Nevertheless, for *k*_d_ = 0.56 s^−1^ and 40 *<* ⟨*n*_b_ ⟩ *<* 70, statistically significant deviations from the 1*/* ⟨*n*_b_ ⟩ scaling law appear, with three data points providing comparable drifting velocities *v*_*y*_. Fig.6b, reporting ⟨*v*_*y*_ ⟩as a function of *k*_d_*/k*_off_ at different ⟨*n*_b_ ⟩, confirms that *k*_d_ has an impact on the drifting velocity for intermediate values of ⟨*n*_*b*_⟩. Overall, Figs. 6a and 6b show how, for strands in which NA molecules are segregated in the tail of the particle (see Fig. 1*b*), the motility of the viruses is controlled primarily by the number of bridges and their interaction range (see next section), followed by the desialysation rate at intermediate values of *n*_b_. It should be stressed that for NA ligands *k*_d_ ≈ *k*_1_ (see Eq. 3) and *k*_1_ is comparable with *k*_on_. Therefore, for ⟨ *n*_b_ ⟩ ≈ 40 the data point in Fig. 6 with the lowest *k*_d_ is the most biologically relevant (see Tab. I).

**FIG. 6.**
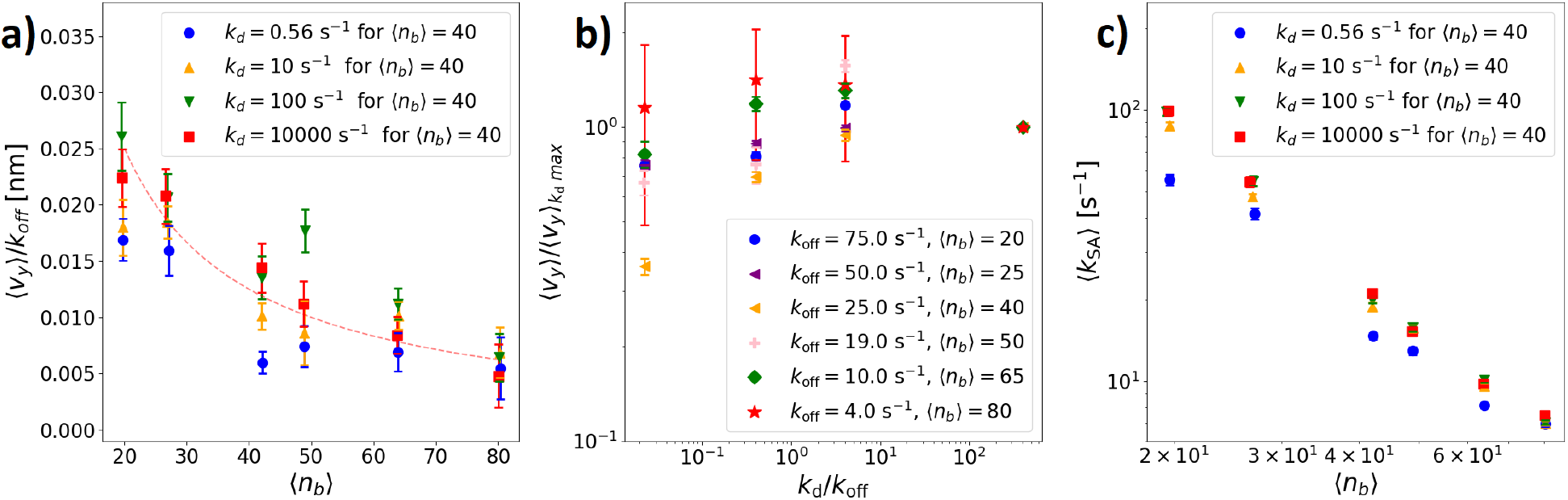
(*a*) Drifting velocity as a function of the number of bridges. We use *k*_on_ = 0.56 s^−1^, change *k*_off_ to tune ⟨*n*_*b*_ ⟩ (see legend of panel *b*), and keep *k*_d_*/k*_off_ constant. The dashed line is a ∼ 1*/*(*n*_*b*_) function fitting the data points with *k*_d_ ≥ 10 s^−1^. (*b*) Drifting velocity as a function of *k*_d_ for the different values of ⟨*n*_*b*_ ⟩ considered in *a*. ⟨*v*_*y*_ ⟩ k_d_max is the mean drifting velocity measured for the largest value of *k*_d_. (*c*) Average rates at which the virus particle inactivates receptors carrying SA as a function of ⟨*n*_*b*_ ⟩. In all panels we use *ρ*_R_ = 0.2 nm^−2^, Δ*t* = 10^−3^ s, and *t*_*E*_ = 100 s. Each data point has been obtained using 30 independent trajectories.

We rationalize the finding of Figs. 6*a* and *b* by sampling the rate at which SA is depleted, *k*_*SA*_ (Fig. 6c). Practically, *k*_*SA*_ is calculated by sampling the number of receptors being desialylated in non-overlapping time intervals of 1 s. Fig. 6c shows how ⟨ *k*_*SA*_ ⟩ is not proportional to *k*_d_. In particular, increasing *k*_d_ leads to a constant value of ⟨ *k*_*SA*_ ⟩ which drastically decreases with ⟨ *n*_b_ ⟩. This trend is explained by the fact that the activity of the NA is reduced by the presence of patches depleted from SA. ⟨ *k*_*SA*_ ⟩ appears to be affected by *k*_d_ for small values of ⟨ *n*_b_ ⟩ while, for the highest values of ⟨ *n*_b_ ⟩, all the *k*_d_ values considered result in a comparable ⟨ *k*_*SA*_ ⟩. With many bridges, the virus moves very slowly, and NA ligands are always able to deplete all the SA in the regions visited by the tail of the particle before moving, no matter the value of *k*_d_. This is peculiar to polarised particles (Fig. 1*b*) in which the NA ligands are segregated and compete to react with the same receptors (see Sec. V for a system in which such a competition is not present). For small values of *n*_b_, particles are faster, and, for small values of *k*_d_, the system cannot cleave the maximal amount of SA. We then interpret the deviation from the 1*/*⟨ *n*_b_ ⟩ scaling law observed in Fig. 6a for *k*_d_ = 0.56 s^−1^ as due to a non optimal desialysation process. The deviation from the 1*/*⟨ *n*_*b*_ ⟩ scaling law is less evident at low values of *n*_b_ (⟨ *n*_b_ ⟩ = 20, 25, Fig. 6a) likely because in these cases we used higher values of *k*_d_ (given that in our simulations *k*_d_*/k*_off_ is constant and that smaller values of ⟨ *n*_b_ ⟩ are obtained by increasing *k*_off_). This is shown in Fig. 6c where it is reported that ⟨ *k*_*SA*_ ⟩ decreases with ⟨ *n*_b_ ⟩ in all cases.

In ESI Fig. 1 we report *k*_SA_ as a function of the drifting velocity of all the systems considered in Fig. 8. Intriguingly, the data points collapse onto a single master curve, showing how *k*_SA_ fully determines ⟨ *v*_*y*_ ⟩. This result also neatly links ⟨ *v*_*y*_ ⟩ to a non-equilibrium parameter.

### B. Interaction range *λ*

In this section, we study the effect of *λ* (defined as the maximal lateral distance between a pair of molecules forming a linkage, see Figs. 1*d*) on the drifting velocity. At a given *k*_on_ and *k*_off_, increasing *λ* would also increase the mean number of bridges, ⟨ *n*_b_ ⟩, because of a multivalent effect. Given that ⟨ *n*_b_ ⟩ and *k*_off_ have a major effect on *v*_*y*_ (Fig. 6a), we designed simulations with three different values of *λ* (*λ* = 3.75 nm, 7.5 nm, and 15 nm) and *k*_on_ (respectively *k*_on_ = 2 s^−1^, 0.56 s^−1^, and 0.12 s^−1^) resulting in a number of bridges equal to ⟨ *n*_b_ ⟩ ≈ 40 (if *k*_off_ = 25.0 s^−1^).

Fig. 7 reports ⟨ *v*_*y*_ ⟩ for three different values of *λ* and two values of *k*_d_. Strikingly, the interaction range has a major impact on the motility of the particle. The increased motility arises from an extension of the border of the surface depleted from SA (see Fig. 5a). The HA ligands close to this region have fewer possibilities to form bridges. The particle then moves away from the SA-depleted region because of a multivalent free energy tempting to maximize the number of bridges (see also Sec. V). The three data points of Fig. 7 predict an increase of ⟨ *v*_*y*_ ⟩ which is proportional to *λ*^3^ (as compared to the 1*/*⟨ *n*_b_ ⟩ scaling law discussed in Fig. 6). This observation points to the fact that maximal motility is achieved not only through fine–tuning of biochemical parameters (like the reaction rates) but also polymeric properties.

**FIG. 7.**
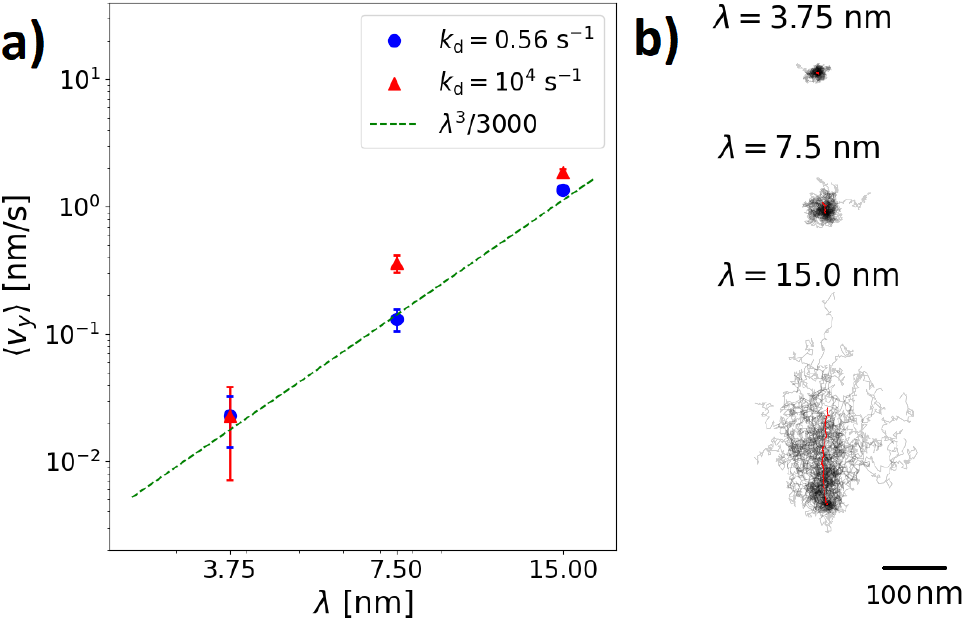
(*a*-*b*) Drifting velocity as a function of the interaction range for two values of *k*_d_. The dashed line shows that ⟨ *v*_*y*_ ⟩ increases like *λ*^3^. We use *k*_on_ [*s*^−1^] = {2.0, 0.56, 0.12}, respectively, for *λ* [*nm*] = {3.75, 7.5, 15} resulting in ⟨ *n*_*b*_ ⟩ = 40. In all cases, *ρ*_R_ = 0.2 nm^−2^, Δ*t* = 10^−3^ s, and *t*_*E*_ = 100 s. Each data point has been obtained using 30 independent trajectories which are reported in panel *b* (the average trajectory is highlighted in red).

Fig. 7 also shows how the activity of the NA ligands, *k*_d_, has a minor effect on ⟨ *v*_*y*_ ⟩ as compared to *λ*. Remarkably, the difference between the two values of ⟨ *v*_*y*_ ⟩ for *λ* = 7.5 nm (discussed in Fig. 6) reduces when decreasing *λ*. In Sec. IV A we argued that, for *λ* = 7.5 nm and ⟨ *n*_b_ ⟩ = 40, low values of *k*_d_ decreases ⟨ *v*_*y*_ ⟩ because the depletion of SA is not maximal (see Figs. 6a and 6c). Instead, Fig. 7 shows how, for *λ* = 3.75 nm, the drifting velocities calculated for the two extreme values of *k*_d_ are comparable (Fig. 7). This result can be interpreted by the fact that, for smaller values of *λ*, each NA ligand can potentially interact with fewer receptors, and maximal depletion is achieved also when using *k*_d_ = 0.56 s^−1^.

### C. Mobile receptors

In this section, we consider receptors diffusing along the cell surface with a diffusion constant equal to *D*_R_ (see Tab. I). In this case, in the chart flow of Sec. III, at each step of the algorithm we also update the position of each receptor *i* by *δ***r**_*i*_ = 2*D*_R_*ξ*_*i*_, where *ξ*_*i*_ is a vector of two independent random numbers following the standard normal distribution. If a receptor is forming a bridge, then the diffusion is limited to a circle of radius *λ* centered over the ligand to which the receptor is bound.

In Fig. 8, we calculate ⟨ *v*_*y*_ ⟩ as a function of *D*_R_ (spanning six orders of magnitudes) for two values of *k*_d_ (Fig. 8*a*) and two different receptor densities (Fig. 8*c*). In these figures, the dashed lines refer to systems with fixed receptors (*D*_R_ = 0). We confirm the trend reported in Ref.^16^, in which the motility is maximized at intermediate values of *D*_R_. The optimal diffusion constant is *D*_R_ = 10 nm^2^*/*s for the lowest value of *k*_d_ (and all receptor densities) and *D*_R_ = 0.1 nm^2^*/*s for the highest value of *k*_d_. Figs. 8*b*, 8*d* show how statistically significant deviations of ⟨ *v*_*y*_ ⟩ from the *D*_R_ = 0 value are concomitant with an increase of the rate at which receptors are inactivated (*k*_SA_). However, further increasing *D*_R_ (and *k*_SA_) does not result in larger values of ⟨ *v*_*y*_ ⟩. Indeed, for large values of *D*_R_, the local gradients generated by the NA ligands are readily lost with the motion of the particles that become diffusive already at the simulation timescales. We note that simulations at high values of *D*_R_ are challenging as they require time steps (Δ*t*) sufficiently small to resolve the motion of a receptor crossing the area scanned by a ligand (corresponding to a circle of radius *λ*, Fig. 3*a*). The results for *k*_d_ = 10^4^ s^−1^ and *D*_R_ ≥ 10^2^nm^−2^ are missing because of a finite-size effect. In particular, in these cases, the NA ligands can inactivate a significant fraction of the receptors present at *t* = 0 (Fig. 5*a*) leading to a significant decrease in the number of bridges ⟨ *n*_*b*_ ⟩.

**FIG. 8.**
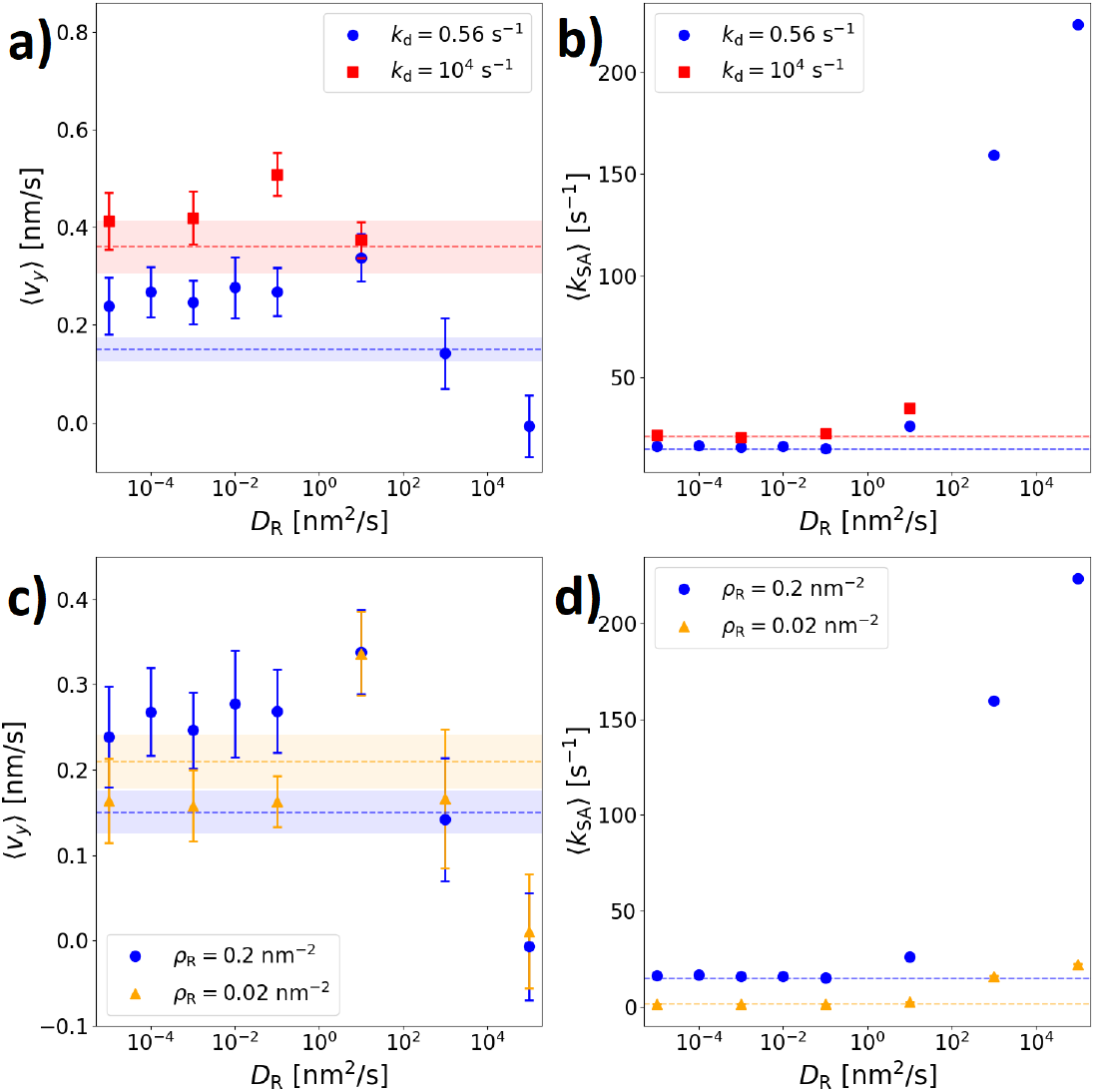
(*a, c*) Drifting velocity as a function of the receptors’ diffusion constant for different values of *k*_d_ (*a*) and receptor density (*c*). (*b, d*) Averaged rates at which the receptors are inactivated corresponding to the simulations of panels (*a, c*), respectively. We use Δ*t* = 10^−3^s *k*_off_ = 25.0 s^−1^ and *k*_on_[*s*^−1^] = {0.56, 5.6} (for *ρ*_R_[nm^−2^] = {0.2, 0.02}, respectively) resulting in ⟨ *n*_b_ ⟩ = 40. Each data point has been obtained using 30 trajectories.

## V. HA AND NA ORGANIZATION ON THE VIRUS SURFACE

As already recognized by Vahey and Fletcher, the segregation between HA and NA ligands at the surface of the virus primes the motility of the particle.^16^ Nevertheless, the organization of the ligands in filamentous IAV strands features a rich variety of patterns and is not limited to the fully polarized configuration considered in Fig. 1*b*.^4,16^ In this section, we show how different ligand distributions have a qualitative impact on the type of movements of the particle.

In Fig. 9, we study the trajectories of particles in which the NA and SA ligands are uniformly distributed across the particle. Each trajectory uses a different randomization of the position of the ligands. In this design, the particles do not drift in the direction of the **y** axis but along the **x** axis. This behavior arises from multivalent forces driving the particle towards configurations that maximize the number of bridges. This is explained in Fig. 10*c*. For particles with uniformly distributed ligands, the depleted region coincides (on average) with the surface scanned by the entire particle. Given that the particle is elongated, a displacement in the **x** direction allows maximizing the number of new receptors entering the reach of HA ligands, and therefore the number of new bridges. Similar considerations justify the fact that polarized particles preferentially move in the positive direction of the **y** axis. Importantly, our previous contribution^24^ clarified how, in the reaction–limited regime, the emerging diffusivity of particles with an arbitrary shape is isotropic (Fig. 10*a*). Therefore, we do not expect the two types of motion described in Fig. 10*b* 10*c* to be affected by anisotropic friction terms.

**FIG. 9.**
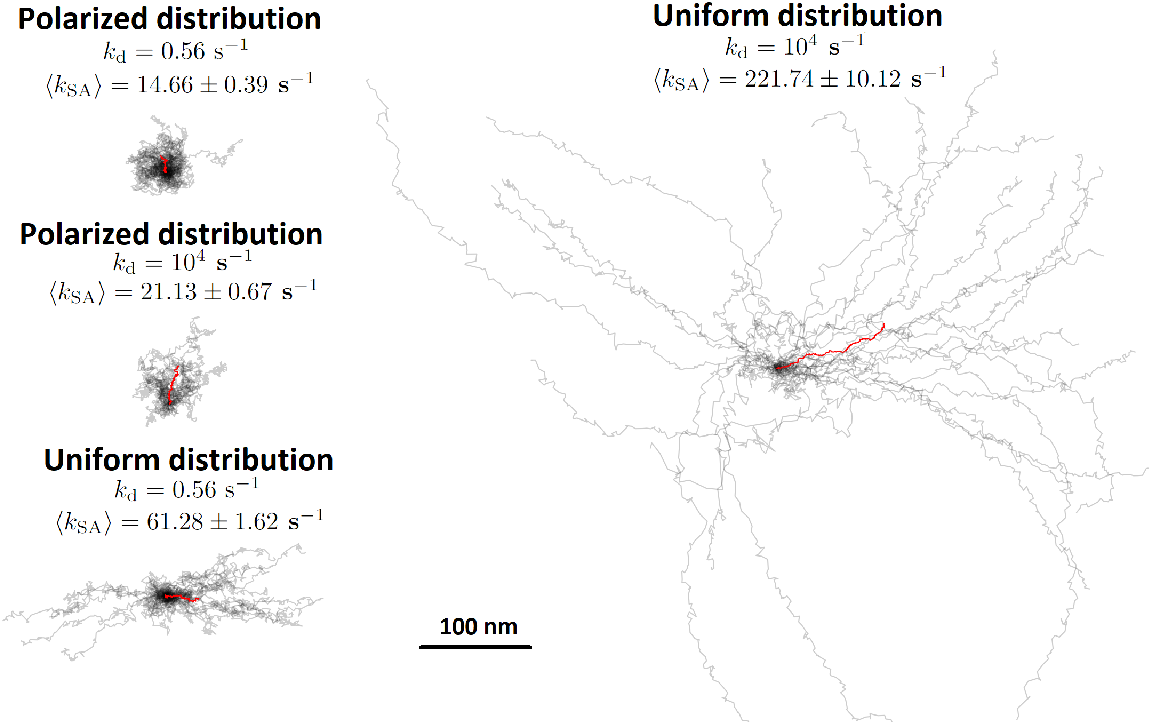
Trajectories of virus particles featuring polarised and uniformly distributed NA ligands. The particle preferentially moves, respectively, along the axial and orthogonal directions. The activity of the NA ligands has a more important role in systems with uniformly distributed NA ligands. We use *k*_off_ = 25.0 s^−1^, *k*_on_ = 0.56 s^−1^, ⟨ *n*_b_ ⟩ = 40, *ρ*_R_ = 0.2 nm^−2^, and Δ*t* = 10^−3^ s.

**FIG. 10.**
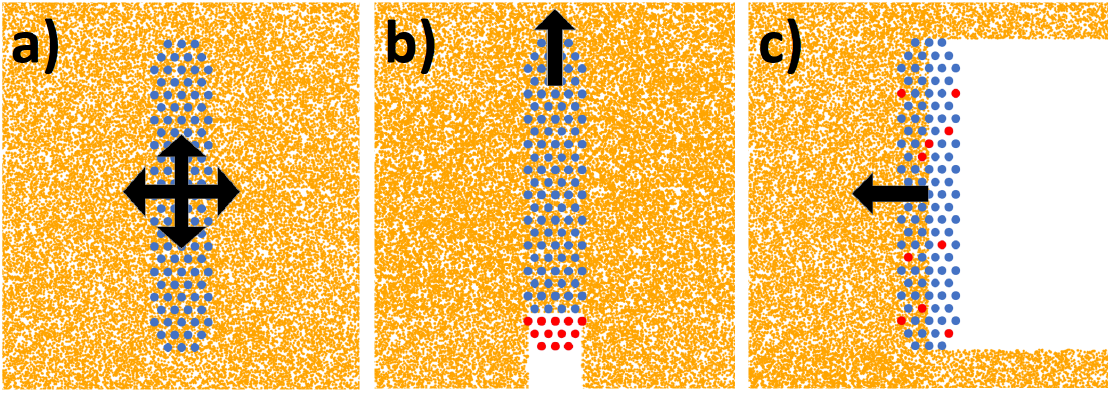
(*a*) In reaction-limited diffusion, without catalytic activity, the distribution of the center of mass trajectories is uniform, irrespective of the particle’s shape.^24^ (*b*) Polarized particles preferentially move along the particle’s axis. (*c*) Particles featuring uniform distributions of NA ligands preferentially move orthogonally to the particle’s axis.

The particle with uniformly distributed ligands does not explicitly break any left/right symmetry. Therefore, starting from the initial configuration without any region depleted from SA, stochastic fluctuations will select an initial random direction and the particle will persist moving in such a direction (Fig. 9). A similar motion has been observed in Influenza C Virus (ICV)^26,50^ and other models implementing burnt–bridge schemes.^27–33^ In ICV, the functional role of binding and cleaving SA is implemented by a single ligand (hemagglutinin–esterasefusion, HEF). In this respect, Fig. 9 shows how the emerging motility of uniformly distributed NA ligands is comparable to ICV systems. We also note that some lateral drifting seems to be present also in polarised particles, at the level of the boundary between HA and NA ligands. For instance, looking at the snapshots in Fig. 5a for *t* = 500 s and *t* = 1000 s, it appears that the particle follows a wiper dynamics, with the tail oscillating left (*t* = 500 s) and right (*t* = 1000 s) (see the movie in the ESI).

Fig. 9 shows how the rate controlling the catalytic activity (*k*_d_) has a more prominent role in the case of particles with a uniform distribution of NA ligands. This arises from the fact that, for uniform distributions, NA ligands can interact with more receptors, and maximal depletion is reached for larger values of *k*_d_. Accordingly, the motility of virions with uniform ligand distributions is much higher than for polarized particles (Fig. 6).

## VI. CONCLUSIONS

Successful infection of Influenza A viruses (IAVs) is based on a synergistic activity between neuraminidase (NA) and hemagglutinin (HA) ligands. While HA ligands anchor the virions to the cell surface by binding receptors carrying sialic acid (SA), NA ligands reduce the number of such NA–HA contacts (or bridges) by cleaving the SA in the intercellular medium. The synergic activity between HA and NA ligands is believed to enhance the motility of the virus particle favoring the spreading of the infection.

In this work, we developed and validated a numerical framework to study the emerging dynamics of IAV particles moving on a surface decorated by receptors carrying SA molecules. We reported on how microscopic parameters featuring a large variability across different IAV subtypes (like the reaction rates) affect the motility of the virion.

The motility of the virus particle (quantified by a short-time, drifting velocity, *v*_*y*_) is mainly controlled by the number of HA–SA bridges (*n*_*b*_) and the interaction range (*λ*), the latter defined as the maximal distance between a pair of interacting ligand/receptor molecules (Fig. 1*d*). In particular, for viruses in which the two types of ligands are segregated (Fig. 10*b*), we found that ⟨ *v*_*y*_ ⟩ ∼ 1*/*⟨ *n*_*b*_ ⟩ (Fig. 5) and ⟨ *v*_*y*_ ⟩ ∼ *λ*^3^ (Fig. 7). Note that the number of bridges is controlled by the *on*/*off* rates (*k*_on_ and *k*_off_, Fig. 1*c*) while the interaction range by the extensibility of the ligand/receptor molecules. Our findings then predict that tiny changes in these parameters have a major impact on the motility of the particles.

We have also studied the effect of the rate controlling the catalytic activity of the NA ligands (*k*_d_) on *v*_*y*_. For segregated ligands (Fig. 10*b*), variations in *k*_d_ have a minor effect (compared to *k*_on_, *k*_on_, and *λ*) on *v*_*y*_ (Fig. 6*a*). In this case, the ensemble of NA ligands can scan a small fraction of the surface (as compared to the particle’s size) with multiple NA ligands potentially interacting with the same receptor. Moreover, the motion of the particle is sufficiently slow to allow the inactivation of all receptors in the reach of a NA ligand, with little difference when considering different values of *k*_d_. This is not the case for uniformly distributed NA ligands (Fig. 10*c*) where different values of *k*_d_ have a major impact on the emerging dynamics and the rate at which receptors are inactivated (Fig. 9).

Finally, different organizations of the ligands on the virus’ surface lead to different types of motion. When considering uniformly distributed NA molecules, the particles preferentially move laterally, while polarized particles slide along their axis (Fig. 9). This observation is general and is driven by the attempt of the particle to maximize the number of bridges *n*_*b*_ (controlling multivalent free-energies^51–53^) along with the fact that, in reaction-limited diffusion, the drag forces (describing the emerging dynamics) are isotropic irrespective of the particle’s shape^24^.

The presence of lateral motility points to the necessity of developing 3D models as, in this case, the rolling component of the motion becomes important.^46^ We anticipate that the computational platform developed in this work could be readily adapted to 3D systems. In the end, this will allow for a quantitative assessment of the motility of virions in extracellular media and, therefore, directly correlate the degree of infectivity of an IAV subtype with the motility of its virions. On the other hand, the results of the present manuscript will inspire the design of complex types of molecular motion and transport.

## Supporting information

Supplemental Fig. 1, supplemental Tab. 1, and caption of the supplemental video

Video supporting manuscript Fig. 5

## CONFLICTS OF INTEREST

There are no conflicts to declare.

## ACKNOWLEDGEMENTS

LS was supported by a seeding grant from the Interuniversity Institute of Bioinformatics in Brussels (IB2). LS and BMM were supported by a PDR grant of the FRS-FNRS (Grant No. T.0158.21). Computational resources were provided by the Consortium des Equipements de Calcul Intensif (CECI) funded by the FRS-FNRS under Grant No. 2.5020.11 and by the Walloon Region.

## Notes

https://github.com/StevensLaurie/IAV

### Competing Interest Statement

The authors have declared no competing interest.

