## Supplemental Fig. 1, supplemental Tab. 1, and caption of the supplemental video for "The sliding motility of the bacilliform virions of Influenza A Viruses"

(Dated:)

[a)] Electronic mail:

### I. SI FIGURE AND TABLE

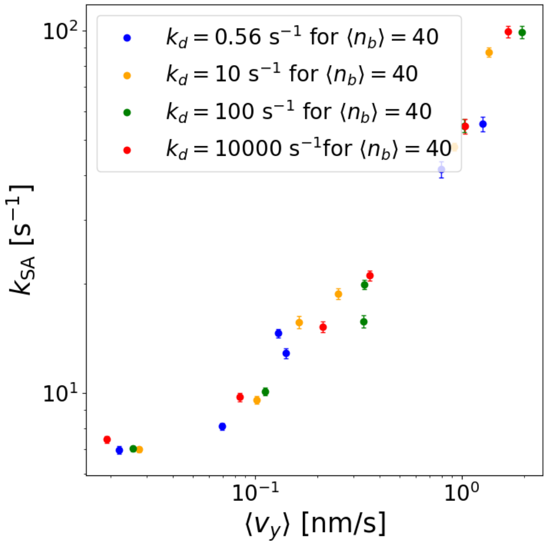


FIG. 1. The rate at which the virus particle cleaves the SA from the receptors determines the drifting velocity. The data points refer to the systems studied in Main Fig. 6.

TABLE I. Effect of the simulation time step ($\Delta t$) on the drifting velocity ($\langle v_y \rangle$). We find that different simulation steps ($\Delta t$) provide consistent drifting velocities, $|\langle v_y \rangle_{\Delta t = 10^{-2}\mathrm{s}} - \langle v_y \rangle_{\Delta t = 10^{-3}\mathrm{s}}| \leq 1.34\,\sigma$. We use $k_{\mathrm{on}} = 0.56\,\mathrm{s}^{-1}$, $k_{\mathrm{off}} = 25.0\,\mathrm{s}^{-1}$ resulting in $\langle n_b \rangle = 40$. The averages have been calculated using 30 trajectories.

| $\Delta t$ | $k_{\mathrm{d}}$ | $\langle v_{\mathrm{drift}} \rangle$ | **Error** $\langle v_{\mathrm{drift}} \rangle$ |
|---|---|---|---|
| | | **Fixed receptors** | |
| $10^{-2}$ s | 0.56 $\mathrm{s}^{-1}$ | 0.205 nm/s | 0.046 nm/s |
| $10^{-2}$ s | $10^4$ $\mathrm{s}^{-1}$ | 0.326 nm/s | 0.066 nm/s |
| $10^{-3}$ s | 0.56 $\mathrm{s}^{-1}$ | 0.154 nm/s | 0.025 nm/s |
| $10^{-3}$ s | $10^4$ $\mathrm{s}^{-1}$ | 0.365 nm/s | 0.054 nm/s |
| | | **Mobile Receptors** | |
| $10^{-2}$ s | 0.56 $\mathrm{s}^{-1}$ | 0.236 nm/s | 0.059 nm/s |
| $10^{-2}$ s | $10^4$ $\mathrm{s}^{-1}$ | 0.334 nm/s | 0.055 nm/s |
| $10^{-3}$ s | 0.56 $\mathrm{s}^{-1}$ | 0.338 nm/s | 0.049 nm/s |
| $10^{-3}$ s | $10^4$ $\mathrm{s}^{-1}$ | 0.375 nm/s | 0.037 nm/s |

#### II. SUPPLEMENTAL MOVIE CAPTION

We report the full trajectory corresponding to Main Fig. 5a.
