## Supplementary figures and images for "The sliding motility of the bacilliform virions of Influenza A Viruses"

### Video supporting manuscript Fig. 5

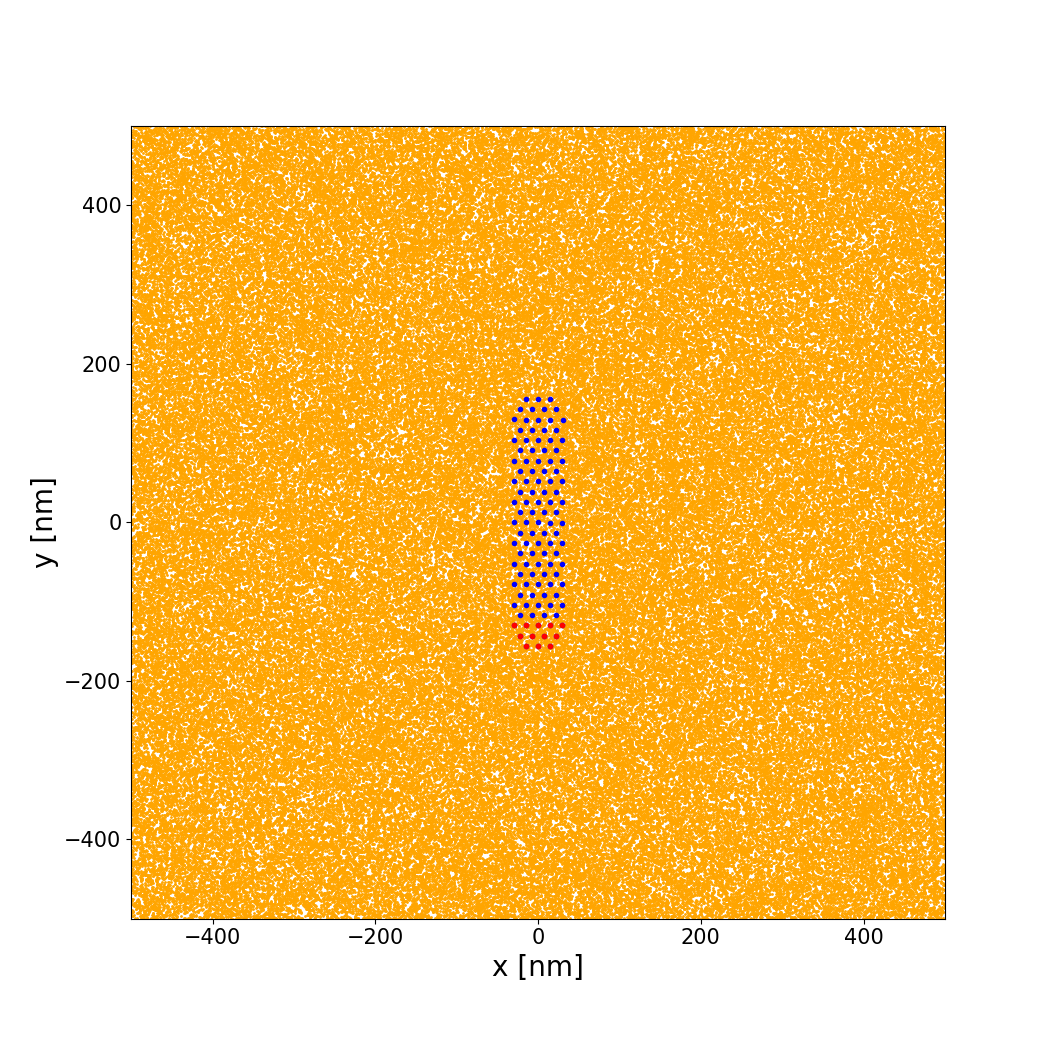
